# Type I IFN stimulates lymph node stromal cells from adult and old mice during a West Nile virus infection

**DOI:** 10.1101/2023.01.05.522898

**Authors:** Allison K. Bennett, Michelle Richner, Madeline D. Mun, Justin M. Richner

## Abstract

Advanced age is a significant risk factor during viral infection due to an age-associated decline in the immune response. Older individuals are especially susceptible to severe neuroinvasive disease after West Nile virus (WNV) infection. Previous studies have characterized age-associated defects in hematopoietic immune cells during WNV infection that culminate in diminished antiviral immunity. Situated amongst immune cells in the draining lymph node (DLN) are structural networks of nonhematopoietic lymph node stromal cells (LNSCs). LNSCs are comprised of numerous, diverse subsets, with critical roles in the coordination of robust immune responses. The contributions of LNSCs to WNV immunity and immune senescence are unclear. Here, we examine LNSC responses to WNV within adult and old DLNs. Acute WNV infection triggered cellular infiltration and LNSC expansion in adult. Comparatively, aged DLNs exhibited diminished leukocyte accumulation, delayed LNSC expansion, and altered fibroblast and endothelial cell subset composition, signified by fewer LECs. We established an *ex vivo* culture system to probe LNSC function. Adult and old LNSCs both recognized an ongoing viral infection primarily through type I IFN signaling. Gene expression signatures were similar between adult and old LNSCs. Aged LNSCs were found to constitutively upregulate immediate early response genes. Collectively, these data suggest LNSCs uniquely respond to WNV infection. We are the first to report age-associated differences in LNSCs on the population- and gene expression-level during WNV infection. These changes may compromise antiviral immunity, leading to increased WNV disease in older individuals.

## INTRODUCTION

West Nile virus (WNV) is a positive sense RNA virus that causes seasonal outbreaks across several continents and continues to threaten public health. WNV is the leading cause of mosquito-borne disease in the United States, however most infections are asymptomatic (McDonald et al., 2021). In some cases (<1%), WNV can infect the central nervous system (CNS), leading to severe neuroinvasive disease and encephalitis (McDonald et al., 2021). Following the bite of an infected mosquito, WNV enters the draining lymph node (DLN), from which it replicates, enters the blood stream, and disseminates into peripheral organs and, eventually reaches the CNS (Suthar et al., 2013). Early innate cues induce inflammatory cytokines that coordinate robust adaptive immune responses, all of which are critical for the containment of WNV disease (Errett et al., 2013; Lazear et al., 2013; Samuel & Diamond, 2005). This coordination is centralized in the DLN where hematopoietic immune cells interact with neighboring stromal cells in lymphoid tissue to facilitate rapid yet controlled adaptive immune responses.

Lymph node stromal cells (LNSCs) constitute less than 2% of the lymph node, and consist of individual subsets that localize to discrete niches of the DLN (Fletcher et al., 2011; Krishnamurty & Turley, 2020). The major well-defined LNSC subsets include fibroblastic reticular cells (FRCs), blood endothelial cells (BECs), and lymphatic endothelial cells (LECs) (Fletcher et al., 2011). Recent single cell RNA sequencing (scRNAseq) studies have identified a number of additional, smaller subsets included within these larger groups (Rodda et al., 2018). LNSCs contribute to anti-viral immunity by integrating inflammatory signals to initiate activation and stimulate adaptive immune responses. Innate signaling pathways, like type I interferon and Toll-like receptors, are stimulated in FRCs and endothelial cell subsets (Gregory et al., 2017; Krishnamurty & Turley, 2020; Malhotra et al., 2012), and an absence of LNSC subsets can lead to attenuated cell-mediated and humoral immune responses during a viral infection (Cremasco et al., 2014). LNSC function can also be disrupted by viral infection, leading to altered lymphoid architecture and, later, impairments in lymphocyte accumulation and organization (Fiorentini et al., 2011; Steele et al., 2009). Infection of FRCs by Ebola (EBOV) and lymphocytic choriomeningitis (LCMV) viruses display shrunken morphology and impaired antigen drainage, respectively (Fiorentini et al., 2011; Steele et al., 2009). However, the mechanism by which LNSCs recognize and respond to an ongoing WNV infection has not been elucidated.

Viral infections, such as SARS-CoV-2, influenza, and WNV, cause increased morbidity and mortality rates in the elderly (>65 years of age) (Bartleson et al., 2021; McDonald et al., 2021).

Furthermore, severe neuroinvasive WNV disease strongly correlates with age in humans, despite similar levels of infection rates across different age groups (Centers for Disease Control and Prevention (CDC), 2019). This disparity is due in part to an age-dependent decline in innate and adaptive immunity known as immune senescence (Nikolich-Žugich, 2018).The mechanisms behind immune senescence are multifactorial. Understanding the mechanisms which lead to this age-dependent susceptibility is critical to develop effective prophylactic and therapeutic strategies.

Animal studies of WNV infection recapitulate the age-dependent decline in antiviral immunity and increased susceptibility. Aged mice infected with WNV display higher viral burdens across multiple organs and increased disease severity (Brien et al., 2009; Funk et al., 2021; Richner et al., 2015). Defects in T cell activation and differentiation, as well as diminished long-lived plasma cell development, have all been attributed to increased disease severity in the aged setting (Brien et al., 2009; Richner et al., 2015). A growing body of evidence suggests that the stromal cells of the lymph node can also contribute to this immune senescence. In both old mice and old non-human primates, the lymphoid tissue microenvironment is disorganized and exhibits a greater level of fibrosis (Kwok et al., 2022; Thompson et al., 2017). Previously, we attributed reduced trafficking of naïve T cells into and within the DLN to defects in the microenvironment of the aged lymph node (Richner et al., 2015). The cellular mechanism leading to these age dependent LNSC defects and how they contribute to the disintegration of the aged immune response, especially during WNV infection, is unclear.

In this study, we examined how LNSCs recognize and respond to an acute WNV infection in both adult and aged settings. Using a mouse model of WNV, we identified age-dependent differences in LNSC population dynamics following an infection. LNSCs from adult and old mice are activated upon WNV infection, primarily through type I IFN signaling. Activation induced similar antiviral gene expression patterns in adult and old LNSC cultures. However, LNSCs from old mice also constitutively upregulated several immediate early response (IER) genes associated with immune suppression. Together these data suggest that LNSCs from adult and old hosts generate an antiviral response to WNV infection through a predominantly IFN-driven mechanism of activation. Age-associated changes in LNSC composition may contribute to the blunted adaptive immune responses and increased severity of WNV infections in older individuals.

## RESULTS

### Aged lymph nodes display alterations in LNSC subsets following WNV infection

To investigate the age-dependent changes in LNSC subsets in a West Nile virus (WNV) infection, we first assessed changes in total cellular dynamics in the draining lymph node (DLN) throughout the acute course of an infection. The rear footpads of adult (8-10 weeks of age) and old (18 months of age) C57BL/6J mice were infected subcutaneously with 1×10^3^ FFU WNV-Kunjin and popliteal DLNs harvested at 2, 4, 6, and 10 days post infection (dpi), or from uninfected mice (0 dpi). In a previous study, we demonstrated that WNV-Kunjin, a less pathogenic variant of WNV, elicits similar lymphocyte trafficking and acute immune responses in the DLN, compared to the more pathogenic strain, WNV-NY99 (Richner et al., 2015). Following inoculation of adult mice with WNV-Kunjin, total cell numbers in the DLN rapidly increased, going from 2.3×10^6^ total cells in the uninfected DLN to 11.9×10^6^ total cells by 2 days post infection (**Fig 1A**). The total number of cells remained elevated in the adult mice at days 2, 4, and 6 post infection before decreasing from day 6 to 10 as infection resolved. On the contrary, old DLNs began with 1.4×10^6^ total cells and increased to only 5.7×10^6^ total cells by 2 days post infection. Total cellularity remained more than 2-fold lower in the old DLN at days 4 and 6 post infection.

**Figure 1:**
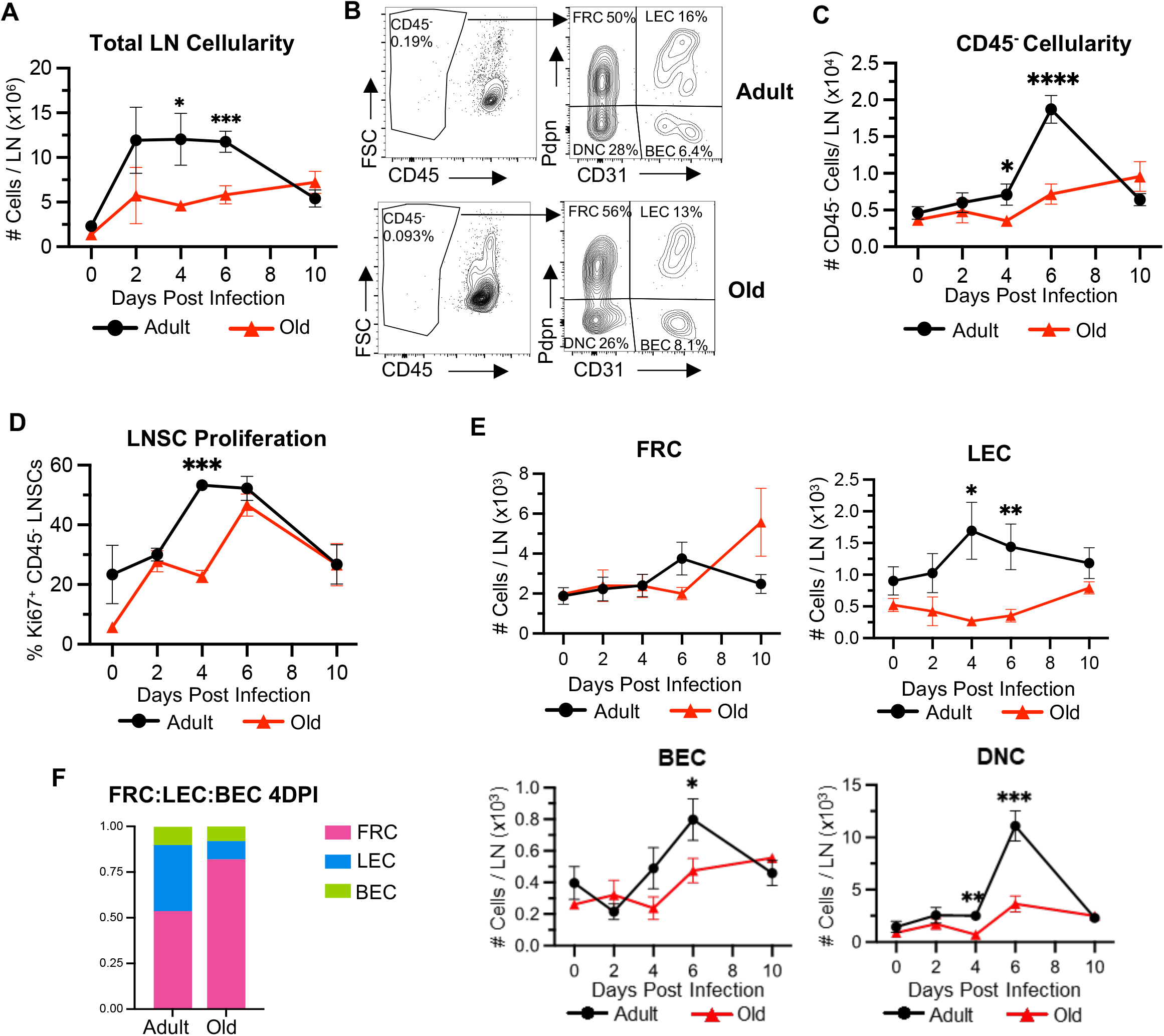
Stromal cell dynamics are altered in aged lymph nodes following WNV infection. Adult (8-10 weeks) and old (18 months) C57BL/6J mice were subcutaneously infected with 1×10^3^ FFU of WNV. Draining popliteal LNs were harvested from infected mice at days 2,4,6, and 10 after infection and digested for flow cytometry. (**A**) Numbers of total cells per DLN. (**B**) Representative dot plots show the gating strategy used to identify LNSCs in the lymph nodes of adult and old mice; percentages on plots represent frequency of cells within the total CD45^-^ population (left plot). CD45^-^ LNSCs were divided into stromal subsets based on Pdpn and CD31 staining (right plot): FRC, LEC, BEC, and DNC. Numbers of (**C)** total CD45^-^ LNSCs and (**E**) FRC, LEC, BEC, and DNC stromal subsets from WNV-infected DLNs of adult and old mice. (**D**) The frequency of proliferating Ki-67^+^ CD45^-^ LNSCs from WNV-infected DLNs were quantified with flow cytometry. (**F**) Ratios of FRC:LEC:BECs from adult and old DLNs at day 4 following WNV infection. The results are averaged from 4 independent experiments with 3-18 mice per timepoint. Data is expressed as the mean ± the standard error of the mean (SEM). Statistically significant differences between adult and old groups at each timepoint is denoted by asterisks (*, *P* < 0.05; ***, *P* < 0.001; ****, *P* < 0.0001; Multiple unpaired T tests).

In addition to measuring total cellularity, we also examined differences in major LNSC subsets in these adult and old infected DLNs. To quantify LNSCs in WNV-infected DLNs, we enzymatically digested popliteal lymph nodes after WNV infection and stained single cell suspensions for CD45, Podoplanin (Pdpn), and CD31 to identify the major LNSC subsets: FRCs, LECs, and BECs. At resting state, the stromal subset found in the highest frequency were the FRCs, followed by LECs and then BECs (**Fig 1B**), similar to the frequencies reported by others (Fletcher et al., 2011; Malhotra et al., 2012). In the adult mice, the numbers of CD45^-^ LNSCs in the DLN peaked at day 6 post infection. LNSC accumulation in the DLN of the old mice had blunted and delayed kinetics, with peak accumulation at 10 dpi (**Fig 1C**). This delayed accumulation of LNSCs in the aged mice correlated with a delayed proliferation as signified by Ki67^+^ staining, a marker for cell division. Adult LNSC proliferation peaked at day 4 and 6 post infection with Ki67+ staining in greater than 50% of CD45^-^ cells. Old LNSCs displayed a delayed pattern of proliferation, peaking at days 6, but was significantly reduced compared to adult at 4 dpi (**Fig 1D**). The dominant proliferating LNSC populations were the FRCs and LECs, with the BECs showing less Ki67 staining over the acute infection course (**Fig S1**). The population of FRCs, BECs, and DNCs in the adult mice, displayed the same pattern of cellular dynamics, with a peak at day 6 (**Fig 1E**). The FRC population in the old mice remained stable from 0-6 days post infection, then increased by 10 dpi. The number of BECs peaked in adult DLNs at 6 dpi, while old LNs displayed a 1.7-fold reduction in this subset. Intriguingly, the number of LECs was consistently lower in the old DLN compared to the adult DLN (p<0.05 at day 4 and 6 post infection). At 4 dpi, old DLNs displayed a significant 6.3-fold reduction in LEC numbers compared to adult (1,694 versus 269, p-value = 0.013). The age-dependent differences in LNSC cellularity are seen more clearly when subset ratios, out of total LNSCs, are compared. FRCs are the dominant subset in both age groups; however, in old DLNs the FRCs comprised a larger percentage of total LNSCs (**Fig 1F**). The LEC compartment is reduced in old DLNs while BECs remain unchanged between both groups. Together these data demonstrate that LNSCs respond and proliferate during an acute WNV infection to support the inflamed, expanding DLN. The expansion of the LNSC network is delayed and blunted in the aged setting.

### LNSCs support WNV replication and are potent producers of inflammatory cytokines

Being the structural network woven throughout the DLN, LNSCs are in the direct presence of viral antigen during an infection. In addition, FRCs and LECs can become infected themselves and serve as a reservoir for viral progeny (Fiorentini et al., 2011; Mueller et al., 2007; Steele et al., 2009). Thus, we first characterized WNV replication in LNSC cultures. To establish lymph node FRC cultures, we isolated and expanded CD45^-^ cells from lymph nodes in the presence of αMEM. These cultures resulted in a purified population of FRC (96.2%) with almost no detectable LEC (0.12%) or BECs (0.024%) (**Fig S2A, C**). LEC and BEC cultures were established from commercially available sources. Individual FRC, LEC, and BEC cultures were infected with WNV-Kunjin (MOI=0.1) for one hour and then washed three times with warm PBS. Supernatants were collected at multiple timepoints following infection and titered by focus forming assay (FFA). Despite multiple washes, virus could be detected from the FRCs at early time points indicating that WNV adheres to the surface of these cells, which did not occur on the BECs nor LECs. In all cell types, viral titers increased over time with a peak titer at 48-72 hpi (**Fig 2A**). These data demonstrate that LNSCs are infected by WNV and can support viral replication.

**Figure 2.**
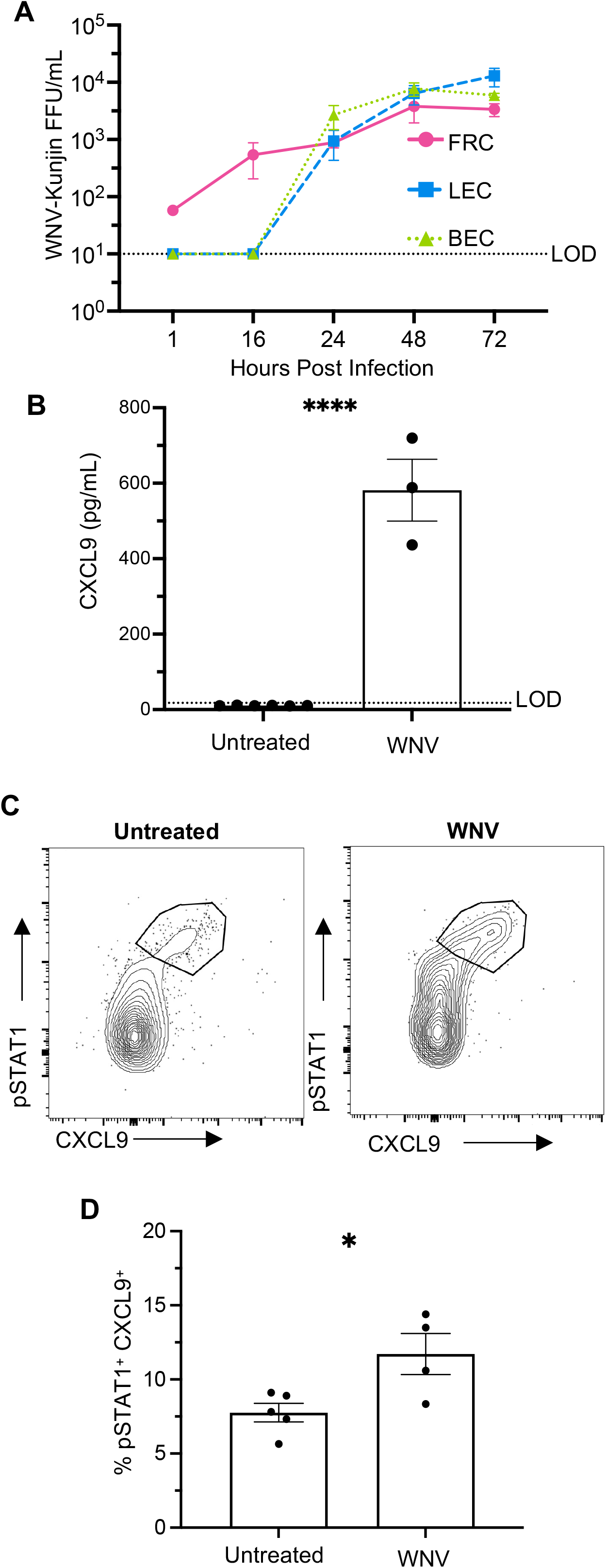
LNSCs support WNV replication and respond to direct infection *ex vivo*. **(A**) FRC, LEC, and BEC monocultures were infected with WNV (MOI= 0.1, 1 hr) and supernatants collected at indicated timepoints. WNV titers were quantified by FFA. The results are averaged from two independent experiments per cell type and data is expressed as the mean titer (FFU/mL) ± SEM. Dashed line represents the limit of detection (LOD= 10 FFU/mL) of assay. (**B**) *Ex vivo* LNSC cultures were infected with WNV (MOI=3) for 20 hours. Cell supernatants were collected to quantify levels of secreted CXCL9 protein via ELISA. (**C-D**) Following infection with WNV, LNSCs were stained with intracellular fluorescent antibodies to CXCL9 and pSTAT1 to quantify stromal cell activation. (**C**) Representative flow cytometry plots of an activated CXCL9^+^ pSTAT1^+^ LNSC population (gated on CD45^-^ cells). (**D**) Frequency of CXCL9^+^ pSTAT1^+^ cells in untreated and WNV treated samples. The results are averaged from two independent experiments with LNSCs from 2-3 mice per experiment. Statistical significance is denoted by asterisks, as compared to untreated (*, *P* < 0.05; ****, *P* < 0.0001; Unpaired T test).

To model LNSC stimulation and define the mechanisms by which LNSCs become activated during WNV infection, we adapted an *in vitro* culture system that contained the major LNSC subsets (Fletcher et al., 2011). Following digestion of pooled skin-draining lymph nodes from adult mice, we plated cells on gelatin coated plates. After 7 days, attached cells were enriched for the presence of LNSCs using CD45 microbeads to magnetically deplete CD45^+^ cells. Following depletion, 95.2% of remaining cells were CD45^-^. Using endothelial cell (EC) media and a gelatin matrix, we grew heterogenous cultures of stromal cells with distinct fibroblast and endothelial cell morphologies (**Fig S2B, C**). Our methods of LNSC enrichment and expansion *ex vivo* resulted in the growth of FRCs, LECs (7.97%), and BECs (4.29%), identified by expression of Pdpn, CD31, and LYVE-1 surface markers. Subsets grown in culture were present in the expected frequencies with FRCs (83.7%) being the dominant cell type. These heterogenous LNSC cultures recapitulated the diverse lymphoid microenvironment *in vivo*, using simple *in vitro* methods (Fletcher et al., 2011; Krishnamurty & Turley, 2020).

We then sought to characterize LNSC activation after a WNV infection. Previous microarray data demonstrated that inflamed FRCs, LECs, and BECs highly expressed *Cxcl9* (Malhotra et al., 2012). CXCL9 is an inflammatory chemokine that drive immune cell migration into and within the DLN and strongly polarize T cells toward an antiviral T_H_1 lineage (Antonelli et al., 2010; Groom et al., 2012). To test *Cxcl9* as an activation readout, LNSC cultures were infected with WNV for 20 hours. CXCL9 was highly secreted by the infected LNSC cultures with a 55-fold (p-value <0.0001) increase compared to unstimulated cells (**Fig 2B**). We further defined LNSC activation through pSTAT1 and CXCL9 intracellular staining in the infected cells (**Fig 2C**). WNV resulted in a 1.5-fold (p-value= 0.026) increase in the percentage of pSTAT1^+^ CXCL9^+^ populations (**Fig 2D**) indicating that WNV can stimulate LNSCs.

### WNV-induced type I IFN activates LNSCs

Within the DLN during infection, LNSCs are exposed to viral antigen and secreted pro-inflammatory cytokines (Krishnamurty & Turley, 2020; Malhotra et al., 2012; Richner et al., 2015). To mimic this inflamed microenvironment, we infected murine bone marrow derived dendritic cells (BMDCs) with WNV-Kunjin (MOI=10) and harvested cell supernatants (“BMDC-WNV”). We characterized the supernatants to determine levels of inflammatory cytokines, IFNα, and infectious virus (**Fig S3**). Stimulation of LNSCs for 20 hours with BMDC-WNV supernatant induced *Cxcl9* expression over 100-fold, as compared to untreated controls (**Fig 3A**). Thus, we have shown that cultured LNSCs are robustly activated upon stimulation with WNV-infected BMDC supernatant, resulting in induction of *Cxcl9*.

**Figure 3:**
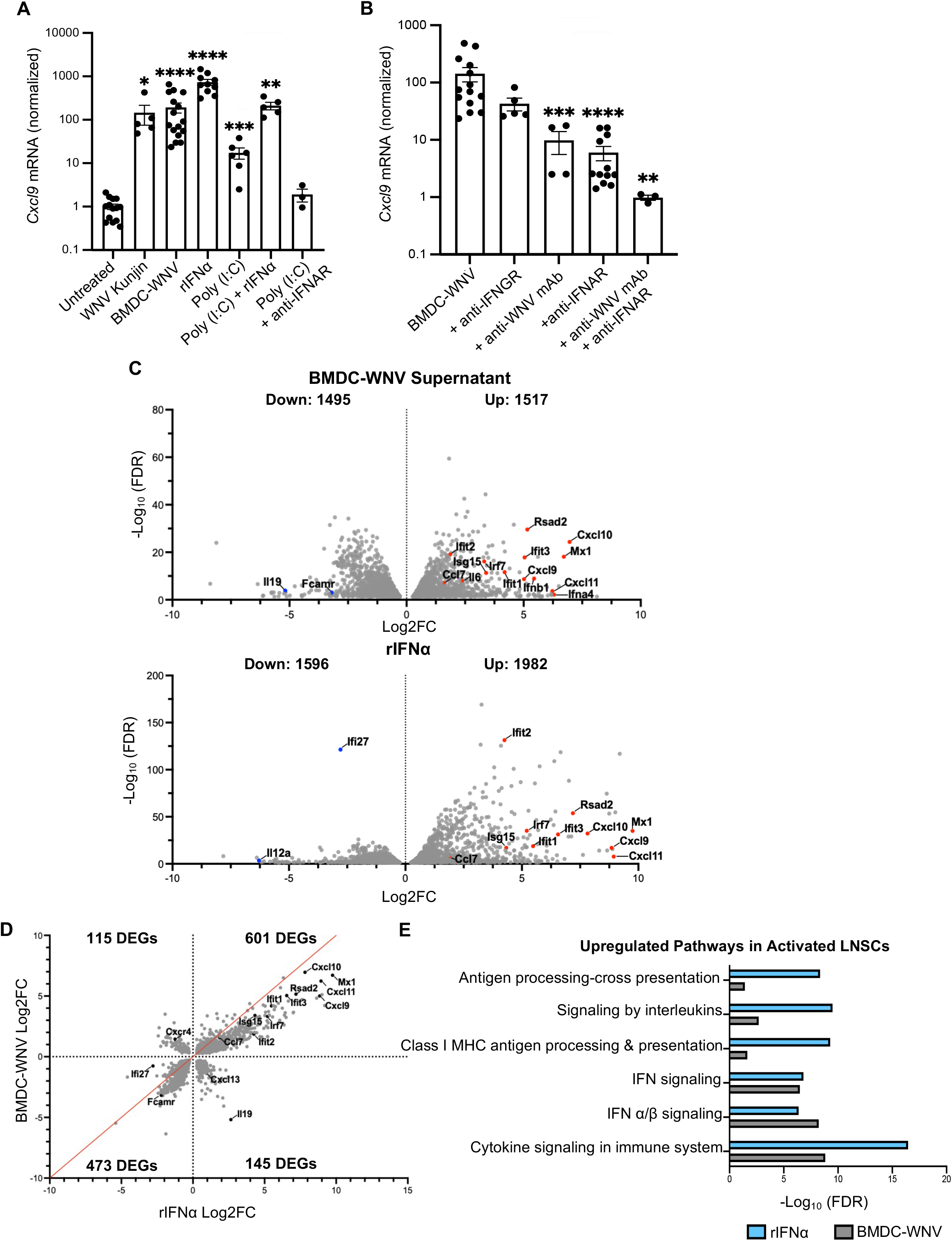
WNV-induced type I IFN activates LNSCs. Ex vivo LNSC cultures from adult mice were incubated with various stimuli for 20 hours and RNA was harvested. (**A**,**B**) Expression *Cxcl9* mRNA was quantified by qRT-PCR. Quantity mean expression levels of *Cxcl9* were divided by *Gapdh* mean expression levels and normalized to untreated. (**A**) Stimulating conditions were: WNV-Kunjin virus (MOI=3), BMDC-WNV supernatant, Poly (I:C) (500 ng/mL), recombinant IFNα (rIFNα, 2 ng/mL), Poly (I:C) + rIFNα, and Poly (I:C) + anti-IFNα receptor (anti-IFNAR, 5 μg/mL). **B**) *Cxcl9* mRNA induction in cultured LNSC after stimulation with BMDC-WNV supernatant and blocking antibody for 20 hours. Antibodies used were: anti-IFN□ receptor (anti-IFNGR, 5 μg/mL), anti-WNV E16 (anti-WNV mAb, 2 μg/mL), anti-IFNAR (5 μg/mL), and anti-WNV mAb + anti-IFNAR. Results are averaged from four independent experiments with LNSCs isolated from 2-3 mice per experiment. Data is expressed as the mean ± SEM. Statistical significance is denoted by asterisks, as compared to untreated (**A**) or BMDC-WNV (**B**) (**, *P* < 0.01; ***, *P* < 0.001; ****, *P* < 0.0001; Mann-Whitney test). (**C**) Global mRNA levels were measure by RNAseq. Volcano plots of BMDC-WNV- or rIFNα-stimulated LNSCs showing differentially expressed genes (DEGs) (FDR< 0.05) compared to untreated cells. Up-regulated (Log2FC >0) and downregulated (Log2FC <0) genes in each comparison are shown, with select genes relevant to WNV infection and the immune response labeled. (**D**) DEGs shared between BMDC-WNV- or rIFNα-stimulated LNSCs were plotted based on Log2FC to visualize changes in magnitude. Line of equivalency is shown in red and number of genes in each quadrant (denoted by the dotted lines) is labeled. (**E**) Functional pathways (REACTOME terms) enriched (FDR< 0.05) in both activated (BMDC-WNV- or rIFNα-stimulated) LNSC groups, as compared to untreated.

We then sought to identify the specific mechanism of LNSC activation observed with BMDC-WNV supernatant. WNV-infected cell supernatants contain high levels of type I IFN and milieu of inflammatory cytokines as well as infectious virus (**Fig S3**) (Lazear et al., 2013; Richner et al., 2015). When mixed cultures of LNSCs were directly infected with WNV virus, *Cxcl9* was upregulated 245-fold (p-value= 0.0167), similar to the level of upregulation with the BMDC-WNV supernatant (**Fig 3A**). Viral PAMPS are known to be potent activators of NF-κB signaling and CXCL9 secretion by triggering type I IFN production (Antonelli et al., 2010; Lazear et al., 2013); however, stimulation with dsRNA analog poly (I:C) (500 ng/ml) resulted in much lower levels of *Cxcl9* induction. Increasing the concentration of poly (I:C) did not augment *Cxcl9* levels (**Fig S4**). When LNSC cultures were stimulated with recombinant IFNα (rIFNα), *Cxcl9* was upregulated (736-fold, p-value= 0.0001) to similar levels as BMDC-WNV supernatant stimulation. Combining rIFNα with poly (I:C) did not induce a synergistic effect. Poly (I:C) induction of *Cxcl9* could be downstream of IFN as RIG-I signaling leads to the upregulation of type I IFN. Indeed, delivery of poly (I:C) with an anti-IFNAR1 antibody, which blocks type I IFN signaling, completely inhibited *Cxcl9* upregulation. These data indicate that type I IFN is a dominant driver of LNSC activation in the context of a viral infection.

To further define the mechanism of LNSC activation, we depleted individual factors from the BMDC-WNV supernatants with monoclonal antibodies. *Cxcl9* can be induced by IFN□ (Antonelli et al., 2010). When LNSC cultures were pre-treated with anti-IFNGR antibody to block type II IFN signaling, and stimulated with infected supernatant, the result was a relatively modest two-fold reduction in *Cxcl9* (p-value= 0.07) compared to BMDC-WNV (**Fig 3A**). Infectious WNV particles in the infected BMDC supernatant might also be driving *Cxcl9*. Neutralization of infectious virus with 2μg/mL of the anti-WNV E16 monoclonal antibody (Nybakken et al., 2005) resulted in a significant reduction of *Cxcl9* up-regulation (100-fold versus 6-fold, p-value= 0.0007) (**Fig 3B**). Similarly, when LNSCs were pre-treated with anti-IFNAR1 antibody prior to BMDC-WNV supernatant treatment, *Cxcl9* was only induced 6-fold, significantly less than BMDC-WNV supernatant (p-value= 0.0001). When WNV particles were neutralized in BMDC supernatant and IFNAR signaling blocked on stromal cells, the result was a complete loss of LNSC activation. Thus, LNSC activation is primarily driven by type I IFN signaling.

We next examined the global gene expression profile of activated LNSCs. RNA was collected from untreated, BMDC-WNV-, and rIFNα-stimulated LNSCs to profile genomic expression using RNA sequencing. BMDC-WNV or rIFNα-stimulated LNSCs were compared to untreated controls to identify differentially expressed genes (DEGs) (false discovery rate, FDR<0.05) (**Table S1**). Both stimulation conditions led to considerable changes in LNSC gene expression (BMDC-WNV 3,012 DEGs; rIFNα 3,578 DEGs), resulting in the identification of upregulated (Log2FC > 0; 1,517 BMDC-WNV, 1,982 rIFNα) and downregulated (Log2FC < 0; 1,495 BMDC-WNV, 1,596 rIFNα) genes (**Table S1, Fig 3C**). Highlighted in both panels of Figure 3C are several upregulated genes in addition to *Cxcl9*, known to mediate aspects of the immune responses and anti-WNV immunity. These genes include *Cxcl9* family members *Cxcl10* and *Cxcl11*, interferon-stimulated genes (ISG) *Rsad2, Isg15, and* antiviral *Ifit1/2/3*. Inflammatory cytokines *Il6* and *Ccl7* were also found to be upregulated across both stimulated LNSCs (**Fig 3C**). BMDC-WNV-stimulated LNSCs were found to downregulate expression of anti-inflammatory cytokine *Il19* and IgA/M receptor *Fcamr*, one of several Fc receptors typically expressed by B cells and some FRC subsets, like follicular dendritic cells (Rodda et al., 2018) (**Fig 3C**). Interestingly, rIFNα-stimulated LNSCs display a downregulation of pro-inflammatory cytokine *Il12a* and ISG *Ifi27*, which has been shown to exhibit antiviral properties during neurotropic viral infection, like WNV (Cho et al., 2013) (**Fig 3C**).

We then assessed DEGs that were up or downregulated (Log2FC >0 or <0) in both stimulation conditions. Changes in magnitude of these DEGs were largely similar across both stimulation conditions (601 upregulated and 473 downregulated DEGs in both groups) (**Fig 3D**). The most strongly shared DEGS, both in identity and change in magnitude, were *Cxcl10, Ifit1, Isg15*, and *Ccl7*. This analysis revealed a smaller group of DEGs whose expression was upregulated in one stimulation condition and downregulated in the other. We found that 115 DEGs upregulated in BMDC-WNV condition and downregulated in the rIFNα condition, this included the chemokine receptor *Cxcr4* that binds the stromal cell-derived chemokine CXCL12 (Cremasco et al., 2014) (**Fig 3D**). In comparison, we found that 145 DEGs were upregulated in rIFNα condition and downregulated in the BMDC-WNV condition. Genes that displayed these expression patterns included *Il19* and stromal cell-secreted chemokine *Cxcl13*, which directs B cell migration within the DLN (Krishnamurty & Turley, 2020). Furthermore, both groups of activated LNSCs showed significant enrichment for the same functional pathways. These included IFN signaling and antigen presentation (**Fig 3E**). This data validates results gathered from earlier *ex vivo* stimulation experiments. Here, we have demonstrated that WNV-Kunjin infection robustly activates heterogenous LNSCs *ex vivo*, leading to the upregulation of ISGs and antiviral cytokines, including *Cxcl9*.

### LNSCs from aged mice have intact IFN response and upregulate immediate early response genes

In addition to the population-level differences, we hypothesize that stromal cell activation is blunted in response to WNV infection of old mice. To investigate differences in the activation of adult versus old LNSCs, we generated *ex vivo* LNSC cultures from the lymph nodes of adult or old naïve mice. Heterogenous LNSC cultures from both adult and old mice grew equivalently, without differences in viability. Adult and old LNSC cultures were stimulated overnight with various stimuli as in previous experiments and *Cxcl9* induction was quantified by qRT-PCR. Adult and old LNSCs displayed similar levels of *Cxcl9* induction after incubation with rIFNα or BMDC-WNV supernatant (**Fig 4A**). The level of activation was reduced in both adult and old LNSC cultures when IFN signaling or WNV particles were blocked. This data revealed that activation of adult and old LNSCs occurred in the same type I IFN-driven mechanism, and these signaling pathways are stable in advanced age.

**Figure 4:**
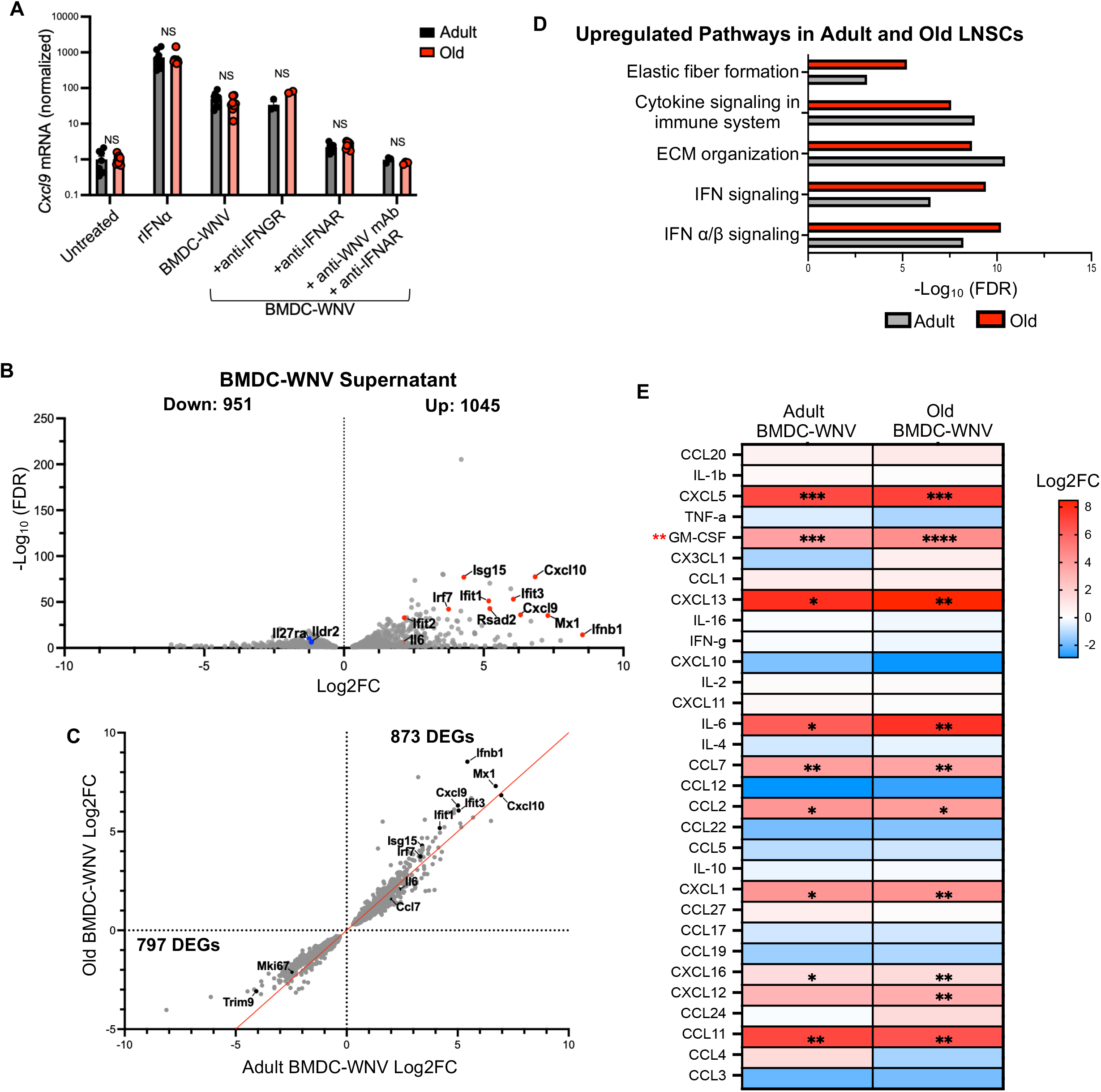
WNV stimuli triggers an antiviral transcriptional response in aged LNSCs *ex vivo*. Ex vivo LNSC cultures from adult and old mice were incubated with various stimuli for 20 hours and RNA was harvested. (**A**) Expression of *Cxcl9* mRNA was quantified by qRT-PCR. The results are averaged from two independent experiments with LNSCs from 4 mice per age group. Data is expressed as the mean ± SEM and expression is normalized to untreated. There were no statistically significant differences (ns, *P* > 0.05; Unpaired t-test). (**B**) Volcano plot of old BMDC-WNV-stimulated LNSCs showing differentially expressed genes (FDR< 0.05) compared to old, untreated LNSCs. Up-regulated (Log2FC >0) and downregulated (Log2FC <0) genes in each comparison are shown, with select genes relevant to WNV infection and immunity labeled. (**C**) DEGs shared between adult and old BMDC-WNV-stimulated LNSCs were plotted based on Log2FC to visualize changes in magnitude. Line of equivalency is shown in red and number of genes in each quadrant (denoted by the dotted lines) is labeled. (**D**) Functional pathways enriched in both BMDC-WNV stimulated adult and old LNSCs, as compared to untreated age-matched controls. (**E**) Supernatants were collected from BMDC-WNV-stimulated adult and old LNSC cultures for quantitative chemokine analysis via 31-Plex Multiplex assay. Heatmap displays Log2FC of indicated chemokines in adult or old LNSC cultures. The results are averaged from two independent experiments. Statistical significance is denoted by asterisks, as compared to BMDC-WNV supernatant alone (black asterisk) or between adult and old LNSCs BMDC-WNV (red asterik) (*, *P* < 0.05; **, *P* < 0.01; Unpaired t test).

We further investigated aged LNSC function by assessing the global gene expression pattern in stimulated aged LNSCs. RNA from adult and old LNSCs that were untreated or stimulated overnight with BMDC-WNV supernatant was collected for RNA sequencing. Using 2-factor multi-group analysis, analogous to 2-way ANOVA, we calculated the effect of stimuli treatment or age on differential gene expression. The interaction term, which captures DEGs where the effect of treatment depended on age, was also calculated. We observed that stimuli treatment significantly drove the variability seen in LNSC gene expression (6,422 DEGs, FDR<0.05) (**Table S2**). Age and treatment did not interact to affect gene expression (0 DEGs, FDR<0.05) (**Fig S5, Table S2**). Activated LNSCs from old mice showed regulation of 1,996 DEGs (1,045 upregulated and 951 downregulated) compared to untreated old LNSCs (**Table S2**). The total number of DEGs in activated, old LNSCs was much lower than those in activated adult LNSCs, as shown in Figure 3C (3,012 DEGs) (**Table S2**). Stimulation of old LNSCs lead to the upregulation of the same antiviral and IFN-stimulated genes highlighted earlier in the adult setting, as compared to untreated old LNSCs. This includes *Cxcl9/10, Ifit1/2/3, Isg15*, and *Il6* (**Fig 4B**). Pro-inflammatory cytokine receptor *Il27ra* and T cell suppressor protein *Ildr2* were found to be downregulated in activated, old LNSCs.

Similar to Figure 3D, we performed further analyses of the DEGs shared between activated adult and old LNSCs and compared their Log2 fold changes in each group. Overall, adult and old LNSCs display strikingly similar patterns of regulation amongst the DEGs they share. Both adult and old, activated LNSCs shared upregulation of 873 DEGs and downregulation of 797 DEGs (**Fig 4C**). Further, the magnitude with which these DEGs changed, especially the ISGs and pro-inflammatory cytokines highlighted, is similar between both adult and old LNSCs as many are located closely to the line of equivalency. Both age groups shared an equivalent downregulation of proliferation marker *Mki67* as well as a downregulation of *Trim9*, a positive regulator of viral-induced type I IFN signaling pathways (Qin et al., 2016). Our analyses did not reveal any DEGs that displayed conflicting patterns of regulation between adult and old stimulated LNSCs. Furthermore, both adult and old LNSCs stimulated with BMDC-WNV supernatant showed enrichment of functional pathways related to IFN- and cytokine-signaling, as well as extracellular matrix remodeling (**Fig 4D**). This further supports our conclusion that IFN responsiveness within old LNSCs remains intact.

After observing changes in pro-inflammatory cytokines and chemokines within both adult and old BMDC-WNV-stimulated LNSCs on the RNA level (Fig 4C), we next wanted to assess changes on the protein level with a 31-plex multiplex cytokine array. We stimulated adult and old LNSC cultures with BMDC-WNV supernatant for 20 hours and collected cellular supernatants for quantification of 31 cytokines/chemokines. Cytokine levels after stimulation were compared to initial cytokine levels in the BMDC-WNV supernatant (**Fig S2**). This analysis revealed significant Log2 fold increases in several chemokines within stimulated adult and old LNSCs (**Fig 4E**). Both age groups exhibited increases in the production of chemokines that organize the migration of immune cells such as B cells (CXCL13), T cells (CXCL16), neutrophils (CXCL1/5), and eosinophils (CCL11), as compared to BMDC-WNV alone (**Fig 4E**, black asterisk). Production of pro-inflammatory cytokine IL6 was also found to be increased in both LNSC cultures. The fold changes in cytokine production between adult and old LNSCs were largely insignificant, except for GM-CSF where old LNSCs produced 1.2-fold more as compared to adult (**Fig 4E**, red asterisk). This data shows that cytokine production and secretion in activated, old LNSCs also remains intact.

To determine if age had any significant impact on LNSC gene expression, we performed pairwise comparisons between untreated adult versus old LNSCs or activated adult versus old LNSCs. These comparisons revealed the age-dependent regulation of 20 and 25 DEGs in untreated and BMDC-WNV-stimulated groups, respectively (**Table S3**). Compared to adult, old LNSCs upregulated 15 DEGs at baseline and 16 DEGs when activated with BMDC-WNV supernatant (**Fig 5A**). Highlighted in Figure 5A, several of these upregulated DEGs were classified as immediate early response (IER) genes and were upregulated regardless of stimulation (Bahrami & Drabløs, 2016). Macrophage markers *Cd209b* and *Marco*, as well as T cell marker *Cd4*, were found to be downregulated the old LNSCs (**Fig 5A**). Some of the upregulated early response genes in old LNSCs, specifically *Nr4a1, Zfp36*, and *Fos*, have been found by others to exhibit immunosuppressive functions (Liu et al., 2019; Moore et al., 2018; Ray et al., 2006). The age associated expression patterns of *Nr4a1, Zfp36*, and *Fos* were validated by qRT-PCR in untreated adult and old LNSCs (**Fig 5B**). We corroborated the expression of ZFP36 in the adult and old infected DLN at 2dpi with WNV-Kunjin. ZFP36 was found predominantly throughout the subcapsular sinus, which would place it in the vicinity of floor and ceiling LECs (**Fig 5C**) (Krishnamurty & Turley, 2020). Overall, LNSCs from aged DLNs are defined by a constitutive, age-dependent upregulation of immediate early response genes, while the IFN-response pathways are unaffected.

**Figure 5:**
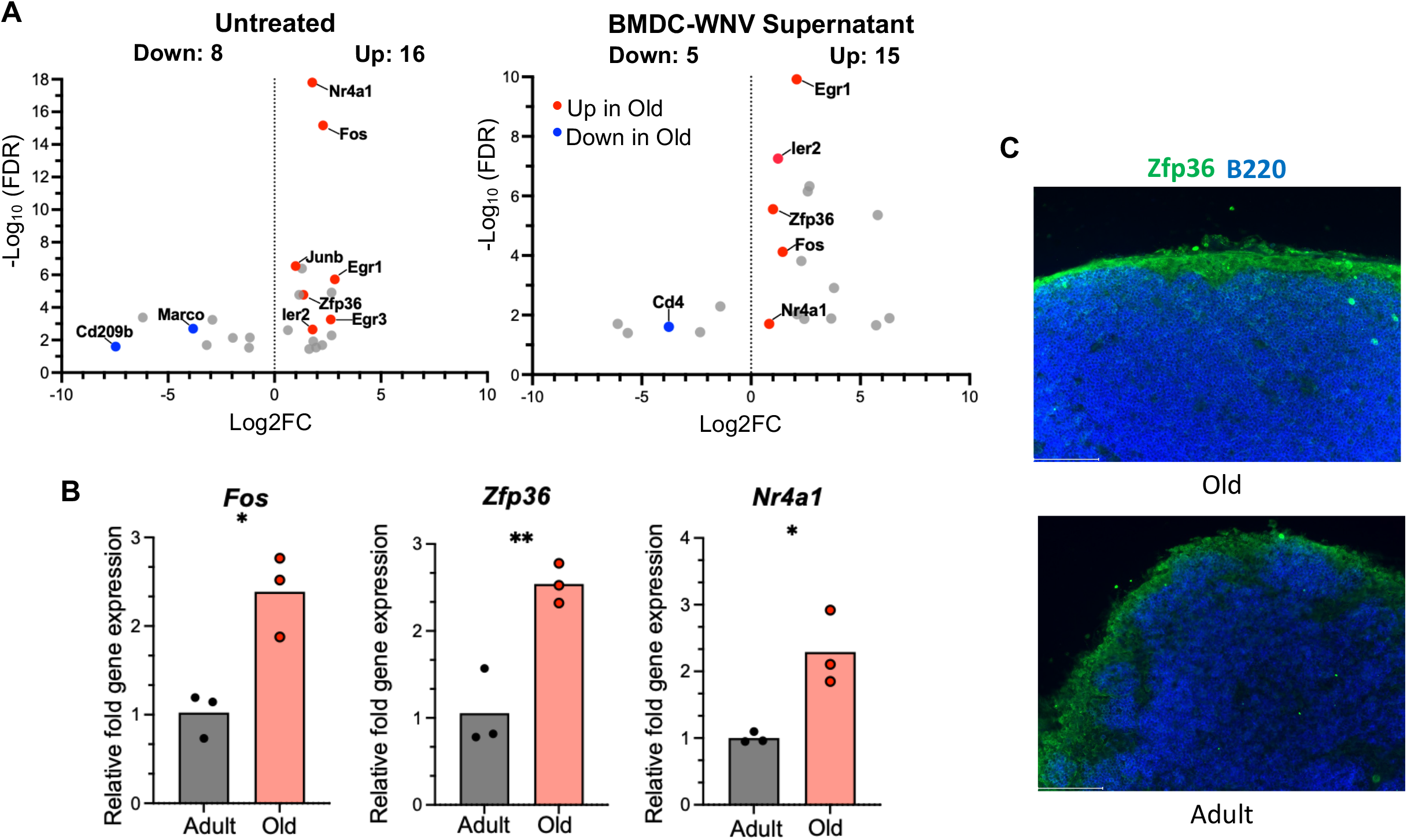
LNSCs display an age-dependent upregulation of immediate early response genes. (**A**) Volcano plot of untreated (left) and BMDC-WNV-stimulated (right) LNSCs from adult and old DLNs comparing differentially expressed genes (FDR< 0.05) between age groups. Up-regulated (Log2FC >0) and downregulated (Log2FC <0) genes in old LNSC cultures, as compared to adult, are shown. Upregulated genes belonging to the immediate early response family are labeled in red and downregulated immune-related genes are labeled in blue. (**B**) Relative fold gene expression of *Nr4a1, Zfp36*, and *Fos* at baseline in adult and old LNSCs, measured by qRT-PCR. The results are averaged from two independent experiments with LNSCs isolated from 2-3 mice per experiment. Data is expressed as the mean ± SEM. Statistical significance is denoted by asterisks, comparing differences between adult and old groups (*, *P* < 0.05; **, *P* < 0.01; Unpaired t test). (**C**) Adult or old mice were infected with 1×10^3^ FFU WNV and draining popliteal lymph nodes were harvested at 2 days post infection. Lymph nodes were cyro-sectioned, and stained for B220 (blue) and ZFP36 (green). Scale bar = 100μm.

## DISCUSSION

Within the DLN resides a diverse population of LNSCs that provide not only structural support, but actively participate in the development of adaptive immune responses. Following an infection of adult mice with WNV, the DLN increased rapidly in size. The rapid increase is due primarily to robust recruitment of naïve B and T cells into the inflamed DLN as we have demonstrated previously (Richner et al., 2015). LNSC levels slightly increased from 0-4 days post infection, but then rapidly expanded from 4-6 days and then contracted by day 10. The increase in LNSCs is due to stromal proliferation (Gregory et al., 2017). Interestingly, the timing and persistence of stromal cell expansion differed between viruses. With influenza, stromal cells displayed peak expansion at 10 DPI and contracted by 12 DPI (Masters et al., 2019). Infection with herpes simplex (HSV) or lymphocytic choriomeningitis virus (LCMV) drove peak stromal expansion by 15 DPI and elevated LNSCs persist beyond day 30 (Gregory et al., 2017). LNSC were productively infected with WNV, similar to findings with other viruses including Ebola virus and LCMV (Fiorentini et al., 2011; Steele et al., 2009). WNV shedding from the BECs could be a mechanism by which WNV reaches the circulatory system after a mosquito bite, leading to systemic viral spread.

Advanced age is correlated with a global decline in immunity, impacting leukocytes and LNSCs. In old mice infected with WNV, we observed reduced leukocyte accumulation and altered stromal subsets. Total cell numbers in old DLNs were significantly reduced at acute time points following WNV, similar to a Chikungunya viral (CHIKV) infection (Uhrlaub et al., 2016). LNSC proliferation was also decreased in the aged DLN. Old WNV-infected mice contained fewer LECs in the DLN during the acute time course. LECs produce CCL21 (Malhotra et al., 2012), therefore fewer LECs could be the cause of diminished CCL21 and reduced T cell trafficking that we have observed previously in the aged setting (Richner et al., 2015). Our results counter findings with influenza virus. Adult and old mice had similar total cell numbers in the lung-draining mediastinal lymph node following an influenza infection, but old mice had reduced numbers of FRCs and LECs (Masters et al., 2019). We postulate that these differences are due to lymph node location as well as pathogen-specific properties. Mucosal antigen exposure within mediastinal lymph nodes throughout life may lead to differences in LNSC priming and expansion, compared to more isolated skin DLNs.

LNSCs are poised to rapidly respond to infection and support the antiviral functions of infiltrating immune cells. Using *ex vivo* heterogenous LNSC cultures, we demonstrated that WNV infection activated LNSCs primarily through a type I IFN-driven mechanism. PAMP activation also led to LNSC stimulation through IFNAR signaling. In agreement, a previous study found that FRCs recognize an ongoing viral infection through type I IFN signaling. The gene expression profiles of the LNSC cultures stimulated with recombinant IFNα or supernatant from WNV-infected BMDCs closely resembled each other. The dominant upregulated pathways were cytokine signaling, and antigen processing and presentation. These data further verify the role of type I IFN in the LNSC response and agrees with previous studies that have analyzed gene expression profiles from individual LNSC subsets (Malhotra et al., 2012). The heterogeneity of our LNSC culture system grown in endothelial cell media recapitulated the DLN microenvironment (Krishnamurty & Turley, 2020), containing expected frequencies of the major LNSC subsets. LNSCs consist of numerous subsets which localize to discrete niches of the DLN, culminating in an exceedingly diverse environment that cannot be perfectly modeled *in vitro* (Krishnamurty & Turley, 2020; Rodda et al., 2018). Therefore, further studies utilizing single cell RNA sequencing will be required to precisely determine virally induced gene expression patterns in the heterogenous LNSCs.

Other findings have identified numerous functional defects in aged LNSCs. Aged LECs suffer from delayed proliferation and increased vessel permeability upon viral challenge (Masters et al., 2019; Zolla et al., 2015). The expansion potential and B/T cell zone architecture of the DLN is shunted due to defects in aged FRCs (Masters et al., 2019; Textor et al., 2016). These aged FRCs formed disrupted reticular cell networks and produced less of T cell chemokines CCL19 and CCL21 following viral infection (Sonar et al., 2022; Masters et al., 2019). Aged BECs are defined by altered expansion kinetics and cellular morphology, the latter of which impedes lymphocyte entry (Masters et al., 2019). To investigate the contribution of aged LNSCs to immune senescence in our model system, we compared adult and old LNSC cultures following stimulation. Counter to our initial hypothesis, LNSCs from adult and old mice robustly induced cytokine expression at equivalent levels. Further, we did not observe any differences in the gene expression signature nor upregulated pathways driven by viral stimulation of the adult and old cultures. Adult and old LNSC cultures also secreted similar levels of cytokines post stimulation. Others have found lower levels of CXCL9 in the aged setting following viral infection. Uhrlaub et al. reported decreased CXCL9 in the serum of old mice with CHIKV (Uhrlaub et al., 2016). Following influenza infection, aged defects in CXCL9 production impaired NK cell accumulation within the DLN (Duan et al., 2017). Lower systemic levels of CXCL9 in the aged setting could be driven by defects found outside of the LNSC compartment.

Despite the overall similarity between the adult and old LNSC cultures, we found that aged LNSCs upregulated immediate early response (IER) genes in both stimulated and unstimulated conditions. Of the upregulated IER genes, *Nr4a1, Zfp36*, and *Fos* are also associated with immune suppression. *Nr4a1* encodes for Nur77, a transcription factor with roles in inflammation and T cell responses (Li et al., 2015; Liu et al., 2019). Old Nur77-deficient mice suffered from systemic inflammation with severe immune cell infiltration and IL6, TNFα production (Li et al., 2015). *Zfp36* encodes for tristetraprolin (TTP), a RNA binding protein that destabilizes target transcripts, especially proinflammatory *Tnf*α and *Il6* (Moore et al., 2018). FOS is an AP-1 transcription factor subunit (Bahrami & Drabløs, 2016), and lack of *Fos* expression *in vitro* and *in vivo* led to increased IL6 and TNFα production following LPS treatment (Ray et al., 2006). Despite the documented roles of these genes in inflammation, these proteins have not been explored in the context of LNSC function nor in the aged immune response. Future studies with conditional overexpression or knock-out mice will be needed to define the role of IER in LNSC function.

In summary, we have characterized the LNSC response to an acute WNV infection and the role for type I IFN-induced activation and PAMP recognition. Overall, LNSCs from aged mice had an intact IFN-dependent response to a WNV infection. A caveat to the current study is that gene-expression data is only monitored from an *ex vivo* culture, and further studies are needed to strengthen our understanding of LNSCs within anti-WNV immunity and immune senescence. Single cell RNA sequencing could allow a more detailed examination of virally induced changes in gene expression on individual LNSC subsets and age-dependent changes in LNSC dynamics. The findings of this study have implications outside the field of viral immunity as advanced age and immune senescence increase the risk of cancer development (Nikolich-Žugich, 2018).

## MATERIALS & METHODS

### Virus and Cells

The WNV-Kunjin strain (CH16532) was generously provided by Michael Diamond at Washington University in St. Louis. Viral stocks were propagated in Vero-E6 cells and titers were determined by focus-forming assay (FFA) as previously described (Brien et al., 2013), using an anti-WNV E16 monoclonal antibody (anti-WNV E16 mAb) (Nybakken et al., 2005). Viral stocks were deep sequenced to validate stock integrity. Experiments utilizing WNV-Kunjin were performed under biosafety level 2 (BSL2) containment at the University of Illinois College of Medicine with approval from the institutional Biosafety committee. Vero-E6 cells (Cat# CRL 1586) were obtained from American Type Culture Collection (ATCC) and maintained in low passage number in accordance with ATCC guidelines. C57BL/6 mouse primary lymphatic endothelial cells (LECs, Cat. C57-6092) and vein endothelial cells (BECs, Cat. C57-6009) were obtained from Cell Biologics. LECs and BECs were maintained using complete mouse endothelial cell medium kit, according to manufacturer’s instructions. Bone marrow-derived dendritic cells (BMDCs) were differentiated in RPMI1640 (Gibco) supplemented with 10% fetal bovine serum (FBS, Gibco), 1% HEPES (Gibco), 1% L-glutamine (Gibco), 1% penicillin-streptomycin (P/S, Gibco), and 20ng/mL GM-CSF (PeproTech) at 37°C for 8 days.

### Mice and WNV Infection

Adult (8-10 weeks) and old (18 months) male C57BL/6J mice were purchased from Jackson Laboratory and housed in pathogen-free conditions at the Biomedical Resources Laboratory at University of Illinois College of Medicine. Mice were infected subcutaneously via footpad injection with 10^3^ FFU of WNV-Kunjin diluted in 50 μl of phosphate-buffered saline (PBS), for a total of 2 × 10^3^ FFU WNV-Kunjin per mouse. Popliteal draining lymph nodes were harvested for further analysis at indicated time points. The WNV infection protocol was approved by the Institutional Animal Care and Use Committee (IACUC) at the University of Illinois College of Medicine (Protocol # 21-084).

### Antibodies

The fluorochrome or biotin-conjugated primary antibodies used for flow cytometry and immunofluorescence microscopy were purchased from Biolegend, eBioscience, Proteintech, or Miltenyi Biotec: CD31 (MEC13.3, 1/200), CD45 (30-F11, 1/200), Pdpn (8.1.1, 1/200), LYVE-1 (ALY7, 1/50), Fc Block (93, 1/200), Ter-119 (1/200), CXCL9 (MIG-2F5.5, 1/200), phospho-STAT1 (A15158B, 1/200), Ki-67 (16A8, 1/100), ZFP36 (12737-1-AP, 1/10), and B220/CD45R (RA3-6B2, 1/10). The fluorescent streptavidin (1/500) secondary antibody was purchased from Biolegend. The anti-WNV E16 mAb used in FFAs and *ex vivo* stimulations was provided by Michael Diamond. The receptor antibodies used in *ex vivo* stimulation experiments: anti-IFN□R (GR-20, Bio-X-Cell) and anti-IFNαR (MAR1-5A3, Leinco).

### Lymph Node Digestion for LNSC Analysis

The LN digestion protocol described below was adapted from a previous protocol (Fletcher et al., 2011). For experiments using cultured LNSCs, inguinal, axial, brachial, and cervical LNs were dissected and pooled in individual digestion reactions per mouse. For experiments analyzing LNSCs following *in vivo* WNV infection, popliteal LNs from infected mice were dissected and each digested individually. Lymph nodes from adult or old mice were dissected and placed into 2mL of basal digestion media comprised of DMEM (Gibco), 2% FBS, 1% P/S, 1% L-glutamine, and 1.2mM CaCl_2_ (Sigma) at room temperature. Once all LNs were dissected, they were punctured once with a 25G needle and placed in tubes containing 2mL of freshly made enzymatic digestion media: basal digestion media, 50mg/mL Collagenase IV (Thermo Fisher), 80mg/mL Dispase (Sigma), and 2000 U/mL DNAse I (Roche). These digestion tubes were placed in a shaking incubator (37°C, 160 RPM) for 25 minutes. After shaking, the LNs were allowed to settle, and the supernatant was moved to a pre-chilled collection tube containing 5mL cell media and 0.5M EDTA. Fresh enzymatic digestion media (2mL) was added to the digestion tube and LNs were gently pipetted until they could pass through a 1mL pipette tip. Digestion tubes were returned to the shaking incubator 10 minutes and the supernatant was transferred to the collection tube after fragments settled, followed by the addition of fresh enzymatic digestion media. LNs were pipetted aggressively until few clumps remained and incubated for 5 minutes. Digestion tubes were removed from the incubator and LNs were pipetted until no clumps remained. All the digested material was added to the chilled collection tube. The contents of the collection tube were passed through a 100μm cell strainer and centrifuged (4°C, 1200 RPM, 5 minutes). LN cells were counted on a hemocytometer, using trypan blue to visually assess cellular viability. Flow cytometry was used to identify LNSCs within LN cell suspensions. Up to 4 × 10^6^ LN cells per sample were stained on ice in FACs buffer (PBS, 1% FBS, .025% NaN_3_) with primary Ter-119, CD45, Pdpn, CD31, LYVE-1, and Fc Block antibodies for 60 min and washed in buffer. Cells were incubated with streptavidin secondary antibody for 20 minutes, washed, and processed on an Attune-NXT flow cytometer. For intracellular cytokine staining, up to 4 × 10^6^ LN cells per sample were stained on ice in FACs buffer with primary surface markers Ter-119, CD45, Pdpn, CD31, and Fc Block antibodies for 60 min and washed in buffer. Cells were then stained on ice in permeabilization wash buffer (0.1% saponin, 0.1% BSA, 1X PBS) with intracellular markers Ki-67, CXCL9, or pSTAT1 overnight. Cells were washed twice in permeabilization wash buffer and processed on an Attune-NXT flow cytometer.

### LNSC Enrichment and Cell Culture

Following digestion, LN cells were resuspended in 10mL cell media and plated on 10cm dishes coated with 0.2% gelatin (Sigma). Complete mouse endothelial Cell Medium (EC, Cell Biologics) was used to grow heterogenous cultures containing FRCs, LECs, and BECs. Minimum Essential Media Alpha (αMEM, Corning) supplemented with 10% FBS and 1% P/S was used to grow homogenous cultures of FRCs. Dishes were placed in a 37°C cell incubator and after 24 hours, washed with PBS and EC media to remove non-adherent cells. EC media was replaced every 3 days. After 7 days, LN cell cultures primarily contained adherent stromal cells with clear fibroblast and endothelial cell morphologies. Stromal cells were incubated with 0.25% trypsin for 2-3 min at 37°C and gently harvested with a cell scrapper. The presence of nonhematopoietic LNSCs was enriched for using CD45 MicroBeads and QuadroMACS™ Separator (Miltenyi Biotec). Up to 10^8^ cells were incubated with CD45 MicroBeads in MACS Buffer (PBS, 0.5% FBS) for 15 minutes at 4°C. The labeled suspension was applied to a LD column on the Separator and the column was rinsed with MACs buffer, according to manufacturer’s instructions. The effluent containing the CD45^-^ cells was collected. Following depletion, 95-99% of the collected cells were CD45^-^. LNSCs were resuspended in EC or αMEM media, counted, and plated on 0.2% gelatin for later experiments.

### Quantification of WNV Infection in LNSCs and BMDC-WNV Supernatants

CD45^-^ LNSCs in EC media were seeded into 24-well plate coated with 0.2% gelatin and incubated in a 37°C incubator. When LNSCs were 80% confluent, WNV-Kunjin was diluted in infection media (DMEM, 2% FBS, 1% HEPES, 1% L-glutamine, and 1% P/S) and added to LNSCs at MOI= 0.1. LNSCs were incubated with WNV at 37°C for 60 minutes, during which the plate was rocked every 15 minutes to evenly disperse virus. Media containing WNV was then removed and LNSCs were washed three times with warm PBS, followed by the addition of fresh EC media. Supernatant was removed from assigned wells at the 1 hour post infection timepoint and the remaining LNSCs were incubated at 37°C. Remaining cell supernatants were collected at 16, 24, 48, and 72 hours post infection and used to quantify WNV infection by FFA, as described previously (Brien et al., 2013). Briefly, serial dilutions of infected LNSC supernatants were incubated with Vero-E6 cells in a 96-well plate for at 37°C for 90 minutes. An overlay containing 1x MEM (Sigma), 1% (w/v) methylcellulose (Sigma), and 2% FBS was then added to Vero cells and incubated. At 32 hours post infection, plates were fixed with 4% PFA. Staining involved primary antibody anti-WNV clone E16 (200ng/mL) and secondary antibody goat anti-human HRP (500μg/mL) in Permwash buffer (PBS, 0.1% Saponin, 0.1% BSA). Incubation with TrueBlue peroxidase substrate (KPL) produced focus forming units that were quantified with the ImmunoSpot® ELISpot plate scanner (Cellular Technology Limited). WNV virus titers in BMDC-WNV supernatants were quantified by FFA as well.

### *Ex Vivo* Stimulation

BMDCs were plated in 12-well plates at 10^5^ cells/well and, following attachment overnight, were infected with WNV-Kunjin (MOI=10) for 20 hours. WNV-infected BMDC supernatants (“BMDC-WNV”) were removed from wells after 20 hours and stored at -80°C. LNSCs were plated in a gelatin-coated 96-well plate with EC or αMEM cell media at a concentration between 5 × 10^3^ and 1 × 10^4^ cells/well and incubated at 37°C overnight. Antibodies were added to LNSCs (anti-IFNAR, anti-IFNGR, anti-LTβR) or BMDC-WNV supernatant (anti-WNV E16 mAb) one hour before stimulation. Plates were returned to the incubator and BMDC-WNV supernatant with anti-WNV mAb was incubated at room temperature. At the time of stimulation, media was removed from all wells and replaced with the assigned stimuli treatment. Cell media (untreated), BMDC-WNV supernatant+/- anti-WNV E16, and WNV virus were added directly to corresponding wells. Recombinant IFNα (Biolegend), TNFα (PeproTech), and IL-1β (PeproTech) as well as poly (I:C) (InvivoGen) were first diluted in cell media and then added to LNSCs. Samples treated with the BMDC-WNV supernatant were infected with an MOI of 5 compared to an MOI of 3 for direct infection with WNV virus, due to the higher concentration of infectious virus in the BMDC supernatant. Anti-IFNAR and IFNGR antibodies were diluted in cell media and added again to antibody treated LNSCs to ensure robust receptor binding. For the anti-WNV mAb + anti-IFNAR condition, anti-IFNAR was added to anti-WNV E16 + BMDC-WNV supernatant and thoroughly mixed. Then, supernatant was applied to anti-IFNAR treated LNSCs. LNSCs were incubated with stimuli at 37°C, followed by the collection of cell supernatant and lysate 20 hours post stimulation. To collect lysate, LNSCs were lysed with Buffer RLT (QIAGEN) and stored at -80°C.

### qRT-PCR

RNA was extracted from LNSC lysates using the RNeasy Mini Kit (QIAGEN). The TaqMan® RNA-to-C_T_™ *1-Step* Kit (Applied Biosystems) was to perform reverse transcription and qPCR in the same reaction. Specific Taqman primers and probes were acquired from Applied Biosystems for *Cxcl9* (Mm00434946_m1), *Nr4a1* (Mm01300401_m1), *Zfp36* (Mm00457144_m1), *Fos* (Mm00487425_m1), and *Gapdh* (Mm99999915_g1). Samples were processed using the ViiA 7 Real-Time PCR system (Applied Biosystems). *Cxcl9* expression was calculated by the division of *Cxcl9* quantity mean expression with quantity mean expression of reference gene *Gapdh*, and then normalized to levels in untreated controls. *Zfp36, Nr4a1*, and *Fos* expression was calculated using the ΔΔC_t_ method.

### RNA-Seq: Sample Preparation, Sequencing, and Bioinformatic Analysis

LNSCs from adult and old C57BL/6J mice were stimulated *ex vivo* overnight with stimuli: BMDC-WNV supernatant, rIFNα (2 ng/mL), or untreated (EC media). Twenty hours post stimulation, RNA was collected, as previously described, and treated with DNase. Each sample was derived from a single mouse and 8 samples per stimuli treatment (N=4 adult LNSCs and N=4 old LNSCs). RNAseq library preparation, quantification, and sequencing were performed by the Genome Research Core at the University of Illinois at Chicago. Briefly, RNA was quantified using the Qubit™ RNA HS Assay Kit (Invitrogen™) on the Quantus Fluorometer (Promega). All RNA samples had a RIN score above 8, as determined by the High Sensitivity RNA Screentape Assay (Agilent) on the Agilent TapeStation 4200 System. RNA samples were normalized to 50 ng and mRNA libraries were constructed using the Universal Plus mRNA-Seq kit (Tecan), followed by RNA-seq on the Illumina NovaSeq6000 S4 flow cell, obtaining 150bp paired-end reads.

Quantification and differential gene expression analysis of RNA-seq data was performed by the Research Informatics Core at the University of Illinois at Chicago. Briefly, raw RNA-seq reads were assessed using FastQC (*Babraham Bioinformatics - FastQC A Quality Control Tool for High Throughput Sequence Data*, n.d.). Filtered reads were aligned to the mouse reference genome mm10 from Ensembl using STAR (Dobin et al., 2013). Genomic features were quantified against Ensembl annotations for mm10 using featureCounts (Liao et al., 2014). Gene expression was normalized to counts per million units and averaged across sample groups, followed by principal component analysis to assess sample clustering. Differential gene analysis was performed in edgeR using 2-factor multi-group analysis to determine effect of treatment and/or age (Robinson et al., 2010). As well, pairwise comparisons were performed between pairs of age and treatment groups, for example “Adult Control versus Old Control” or “Old BMDC-WNV versus Old Control.”

The gene expression profiles from adult and old LNSCs following *ex vivo* stimulation were analyzed for significant enrichment of functional pathways using the ToppFun application within the ToppGene Suite (Chen et al., 2009). Upregulated differentially expressed genes (FDR <0.05) with a log2-fold change greater than 0 were used for pathway enrichment analysis. Results were filtered to only REACTOME pathway terms with FDR <0.05 (Jassal et al., 2020).

### Quantitative Cytokine Analysis

Supernatants from stimulated adult and old LNSC cultures, or BMDC WNV supernatant alone were collected and stored at -80°C. Chemokines were quantified using a Bio-Plex Pro Mouse Chemokine Panel (Bio-Rad). IFNα (Cat. 421201) and CXCL9 (Cat. DY492-05) were quantified by ELISA kits from Invitrogen and R&D Systems, respectively.

### Immunohistochemistry and fluorescence microscopy

Popliteal draining lymph nodes from adult and old mice were harvested 2 days post infection with WNV-Kunjin. DLNs were frozen in O.C.T. blocks and 8-μm-thick sections were cut using a cryostat. DLN sections were mounted on microscope slides, stained with anti-ZFP36 (ProteinTech, Cat.# 127-37-1-AP, 1/10) and anti-B220 (Biolgened, Cat. # 103239, 1/10) fluorescently conjugated antibodies, and imaged with a Keyence BZ-X700 series microscope.

### Data Analysis

All data, excluding bioinformatic analyses, was analyzed using GraphPad Prism 9 software (San Diego, CA). Flow cytometry data was analyzed with FlowJo software (BD Biosciences). Statistical significance was calculated via Mann-Whitney test or unpaired-T test as indicated in Figure legends.

## ACKNOWLEDGEMENTS

This work was funded by a grant from the National Institute of Allergy and Infectious Diseases (R01AI150672) and institutional startup funds to J.M.R. RNA sequencing and basic processing of the raw data were performed by the University of Illinois at Chicago Genome Research Core and Research Informatics Core, respectively. The graphical abstract was generated using BioRender.com.

## AUTHOR CONTRIBUTIONS

A.K.B and J.M.R. designed the experiments and analyzed the data. A.K.B and M.R. performed experiments and A.K.B. generated the figures. M.D.M performed immunofluorescent microscopy experiments. A.K.B and J.M.R wrote and edited the manuscript, with input from M.R. and M.D.M.

## DATA AVAILABILITY

Bulk RNA sequencing datasets generated by this study have been submitted to the NCBI Gene Expression Omnibus (GEO) repository under accession number GSE198629.

## FIGURE LEGENDS

**Supplementary Figure 1:**
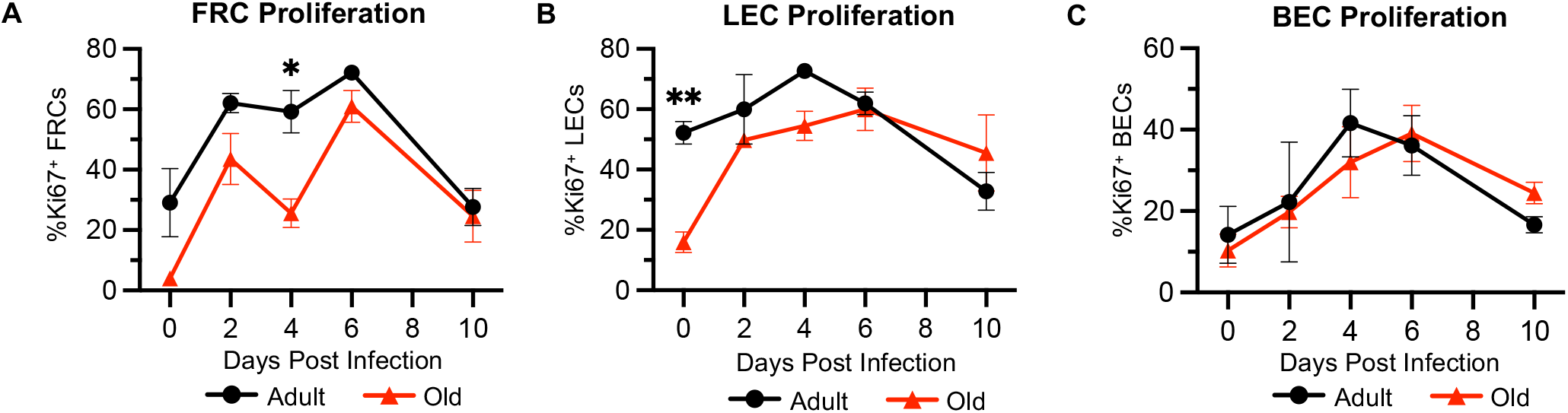
WNV infection triggers expansion of LNSC subsets in adult and aged DLNs *in vivo*. Adult (8-10 weeks) and old (18 months) C57BL/6J mice were subcutaneously infected with 2×10^3^ FFU of WNV-Kunjin. Draining popliteal LNs were harvested from infected mice at days 2,4,6, and 10 after infection. At each timepoint, frequencies of proliferating Ki67+ FRCs (**A**), LECs (**B**), and (**C**) BECs were quantified in adult and old DLNs. The results are averaged from 2 independent experiments with 3 mice per timepoint. Data is expressed as the mean ± SEM. Statistically significant differences between adult and old groups at each timepoint is denoted by asterisks (*, *P* < 0.05; ***, *P* < 0.001; Multiple unpaired T tests).

**Supplementary Figure 2:**
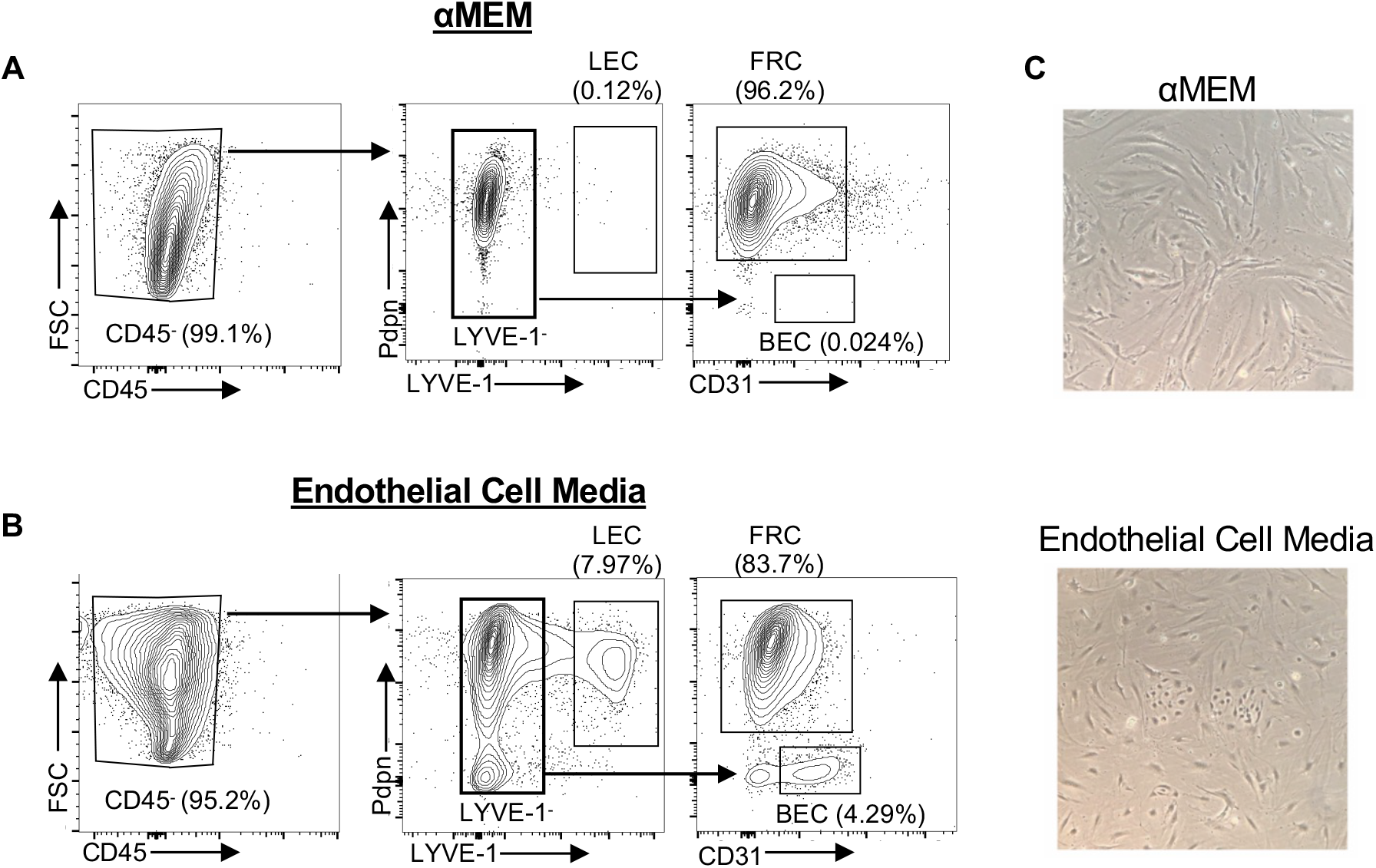
Characterization of *ex vivo* LNSC culture systems. LNSCs from skin draining LNs from adult C57BL/6J mice were digested and expanded in gelatin-coated dishes with (**A**) αMEM media or (**B**) endothelial cell medium. Growth of major LNSC subsets was confirmed by flow cytometry. (**A**,**B**) Representative dot plots show the gating strategy used to identify LNSC subsets in each culture system. LNSCs (gated on CD45^-^ cells) were divided into LEC (LYVE-1^+^) and non-LEC (LYVE-1^-^) populations based on Pdpn and LYVE-1 staining (left). The LYVE-1^-^ population was divided into FRCs and BECs based on Pdpn and CD31 staining (right). (**C**) Representative images of *ex vivo* LNSC cultures grown in the presence of αMEM media (top) or endothelial cell medium (bottom).

**Supplementary Figure 3:**
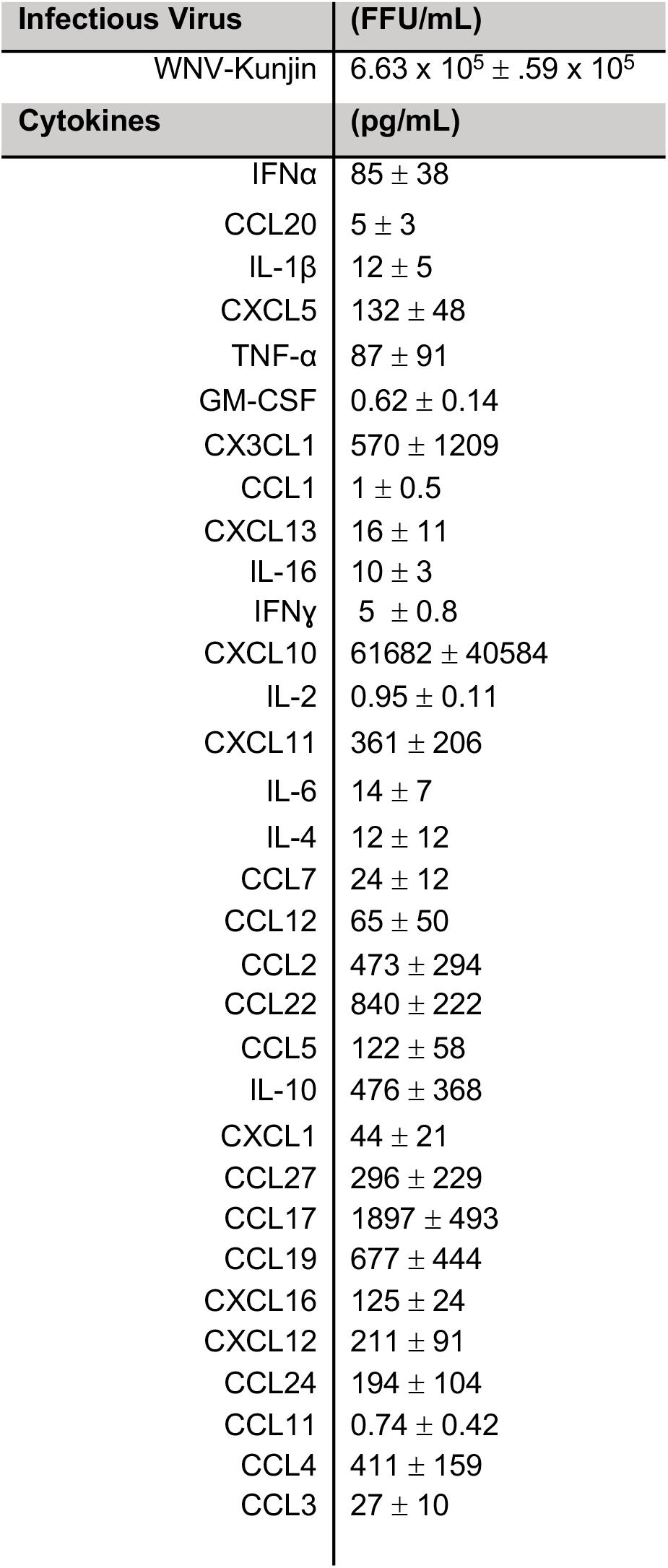
BMDC-WNV supernatants contain both inflammatory cytokines and infectious virus. Bone marrow-derived dendritic cells (BMDCs) were infected for 20 hours with WNV-Kunjin virus (MOI=10), and supernatants were collected for further characterization. Infectious WNV-Kunjin titers were quantified by FFA. Secreted levels of IFNα and chemokines were quantified by ELISA and 31-Plex Multiplex assay, respectively.

**Supplementary Figure 4:**
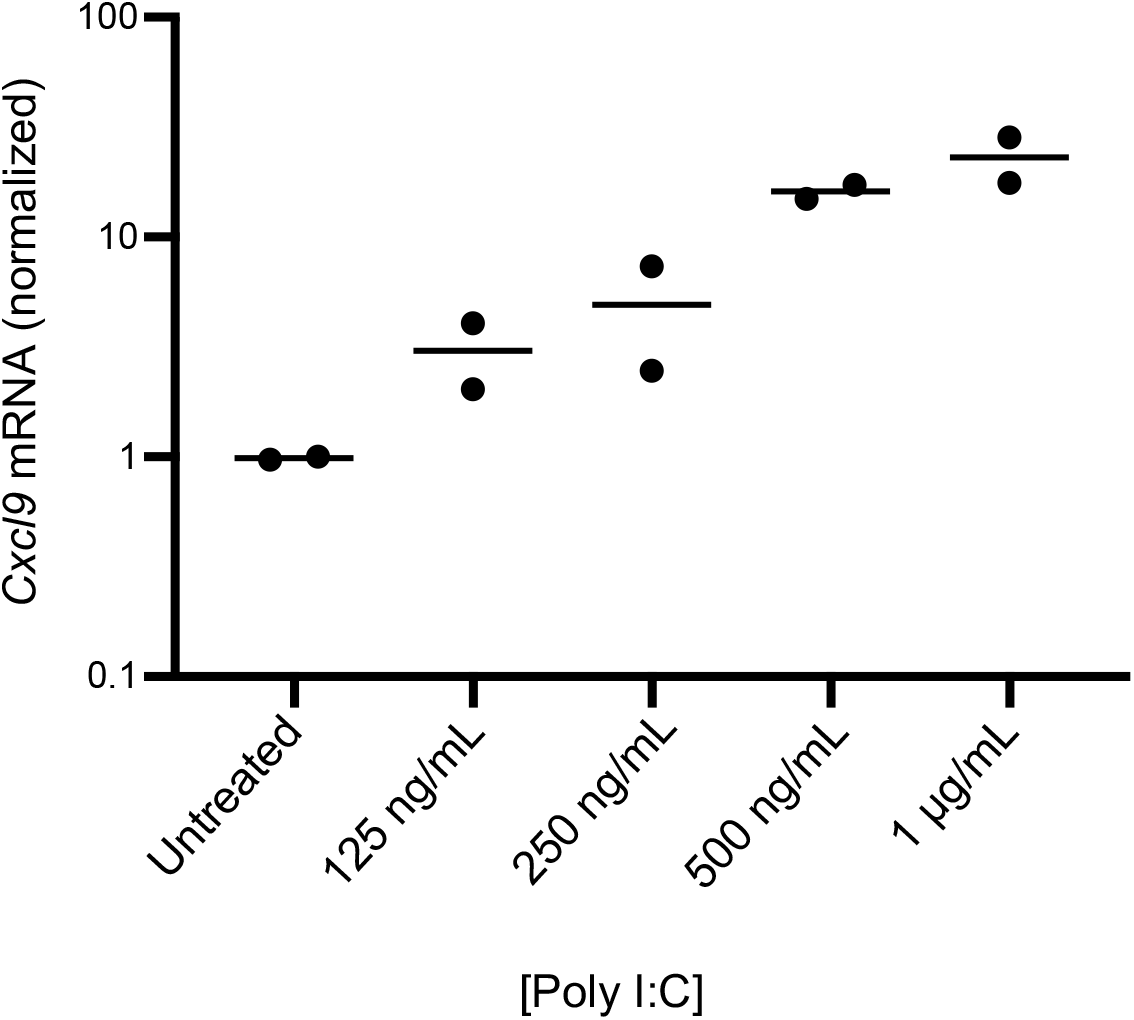
LNSC activation is not augmented by increasing poly I:C concentration. LNSCs cultured in endothelial cell media were incubated with increasing concentrations of poly I:C (125 ng/ml-1μg/mL) for 20 hours. *Cxcl9* expression was quantified by qRT-PCR and normalized to untreated. Data is expressed as the mean ± SEM.

**Supplementary Figure 5:**
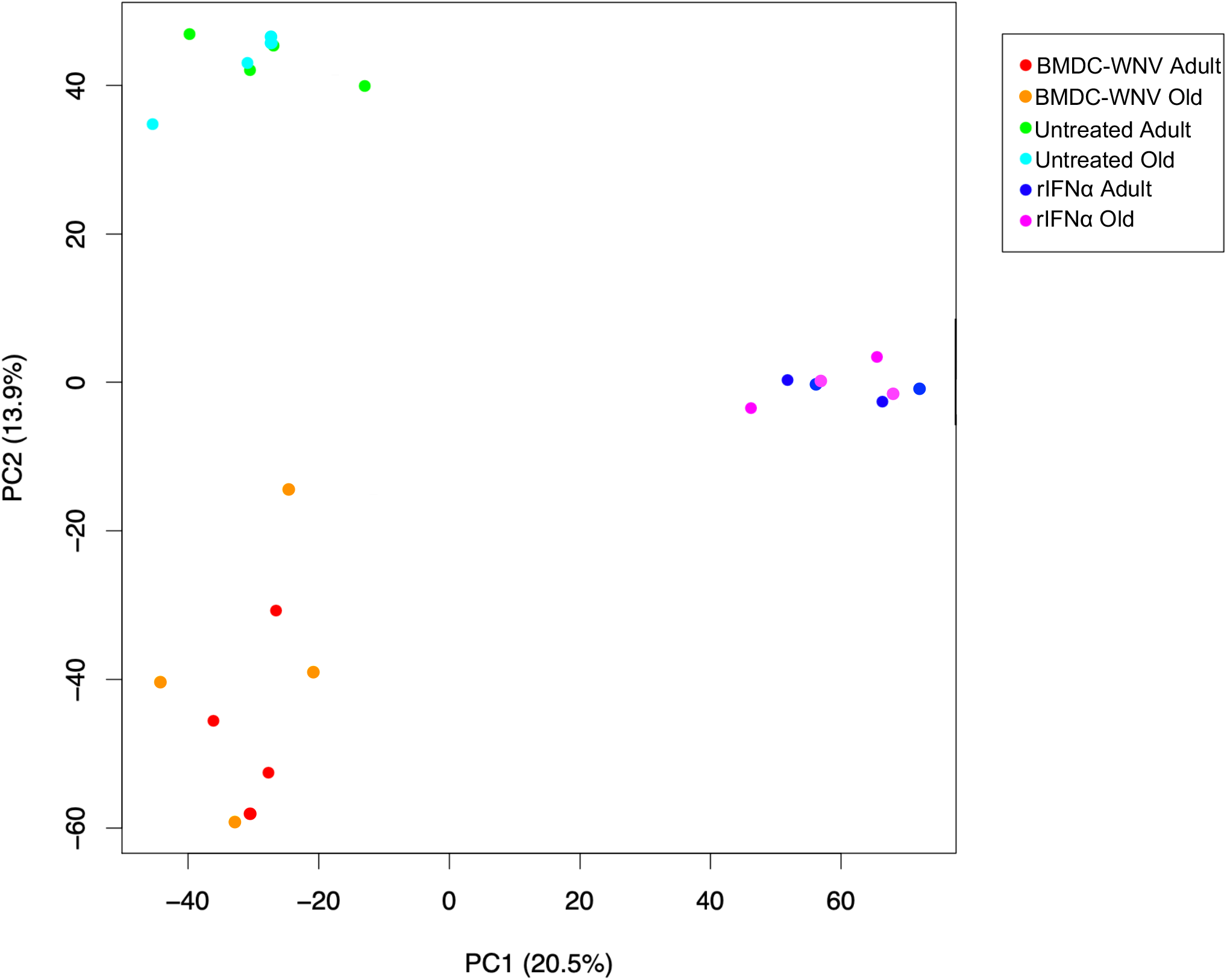
Age does not significantly drive global gene expression variability in LNSCs. Principal component analysis (PCA) was performed on normalized RNA-seq data of adult and old LNSCs from *ex vivo* stimulation experiments: untreated, BMDC-WNV, and rIFNα. Each dot represents a single sample and is colored based on sample type. Adult and old samples cluster together based on treatment, rather than age.

